# Analyzing associations and higher-order effects in multi-omics data with double machine learning

**DOI:** 10.64898/2025.12.04.691622

**Authors:** Julian Hecker, Dmitry Prokopenko, Georg Hahn, Sanghun Lee, Sharon M. Lutz, Christoph Lange

**Affiliations:** Channing Division of Network Medicine, Brigham and Women’s Hospital and Harvard Medical School, Boston, MA, USA; Genetics and Aging Research Unit and McCance Center for Brain Health, Department of Neurology, Massachusetts General Hospital and Harvard Medical School, Boston, MA, USA; Division of Pharmacoepidemiology and Pharmacoeconomics, Brigham and Women’s Hospital and Harvard Medical School, Boston, MA, USA; Department of Biostatistics, Harvard T.H. Chan School of Public Health, Boston, MA, USA; Department of Medical Consilience, Division of Medicine, Graduate School, Dankook University, Yongin-si, South Korea; Department of Population Medicine, Harvard Pilgrim Health Care Institute and Harvard Medical School, Boston, MA, USA

## Abstract

**Motivation:** Integrative omics analyses enhance our understanding of disease mechanisms and biomarkers by investigating relationships among traits, omics measurements, genetic variants, and epidemiological factors. Statistically, these analyses are challenging and require robust, flexible methodologies due to high dimensionality, non-standard data distributions, and potentially complex, non-linear confounding effects.

**Results:** To facilitate the integration and analysis of multi-omics data, we introduce the Robust Omics MethodologY (ROMY) framework and its corresponding R implementation, the *romy* package. ROMY enables users to (a) perform robust association testing between two target variables while incorporating flexible covariate adjustments, (b) examine effects on measurement variances and covariances (e.g., co-expression, co-abundance), and (c) conduct rigorous interaction-effect testing. ROMY builds on recent advances in theoretical statistics and double machine learning to ensure robustness and statistical validity. We illustrate the performance of our framework through simulation studies.

**Availability and implementation:** *romy* is an R package available under the GNU GPLv3 license on GitHub: https://github.com/julianhecker/romy.

## Introduction

Integrative omics analyses aim to unravel the molecular mechanisms underlying complex traits and to identify relevant biomarkers^1,2^. Such analyses examine the relationships between complex traits, omics measurements (including genetics, epigenetics, metabolomics, and proteomics), and epidemiological variables.

Multiple factors increase the complexity of integrative omics analyses: omics data are typically high-dimensional, measurements are often noisy and exhibit non-standard distributions, and (confounding) effects can be scale-dependent and non-linear. These challenges underscore the need for robust methods for omics analysis and their efficient implementation.

Here, we introduce our Robust Omics Methodology (ROMY) framework, which aims to address these challenges and demands. Our framework can a.) investigate the association between a variable of interest (for example, a genetic variant, omics measurement, or epidemiological factor) and a quantitative outcome (omics measurement or complex trait) while robustly adjusting for confounding variables, b.) assess effects on covariances and variances, and c.) perform robust interaction-effect testing. These components are based on the concept of debiased/double machine learning^3,4^ and one-step corrected estimators^5^, enabling us to adjust for confounding or precision variables using modern machine learning models while obtaining valid statistical tests. We conduct simulation studies to demonstrate the validity of the proposed methodology. Our framework is implemented in the R package *romy*, whose modular design allows researchers to integrate any suitable future statistical learning model for covariate/confounder adjustment.

## ROMY framework

The ROMY framework comprises three analysis components: ROMY-CIT, ROMY-COV, and ROMY-INTER. In the following, we introduce and motivate each component. Details of their implementation and theoretical details are provided in the Supplementary Materials.

### Setting

Let the vector *Y* = (*Y*_1_, *…, Y*_*M*_) denote a set of *M* quantitative outcomes (complex traits or omics measurements). Next, let *X* = (*X*_1_, *…, X*_*P*_) denote a different layer of omics or epidemiological factors with *P* components. For example, *Y* could represent metabolites and *X* expression data. Moreover, the *D*-dimensional variable *Z* denotes covariates that the analyses aim to adjust for. We assume that this data is collected across a joint set of samples of size *N*, resulting in observed data *Y*_*i*_ = (*Y*_*i*1_, *…, Y*_*iM*_), *X*_*i*_ = (*X*_*i*1_, *…, X*_*iP*_), and *Z*_*i*_ = (*Z*_*i*1_, *…, Z*_*iD*_), *i* = 1, *…, N*.

### ROMY-CIT: association testing

The first analysis approach focuses on testing for the association between *Y*_*m*_ and *X*_*p*_, *m* ∈{1, *…, M*} and *p* ∈ {1, *…, P*}, while adjusting for the covariates *Z*. Our approach is based on the so-called generalized covariance measure^6^ for conditional independence testing (CIT), given by

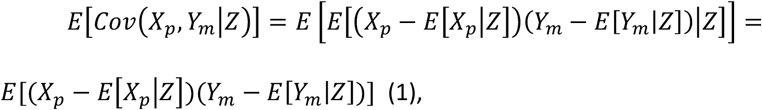

which equals 0 under the null hypothesis that *Y*_*m*_ and *X*_*p*_ are independent given *Z*. This functional, the expected conditional covariance, was also extensively studied in the causal inference literature^3,5,7^. Theoretical results show that it is possible to use machine learning prediction models or other flexible statistical learning algorithms to estimate *E*[*X*_*p*_ *Z*] and *E*[*Y*_*m*_|*Z*] and plug their estimates 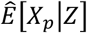 and 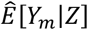 into the test statistic component

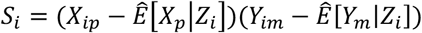

for individual *i*, to obtain valid statistical inference to test *E*[*Cov*(*X*_*p*_, *Y*_*m*_ *Z*)] = *0* based on 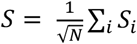 under mild assumptions. This is attributable to the specific product form of the functional that requires only the product of the estimator’s convergence rates to be fast. That is, the rate of 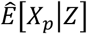 converging to *E*[*X*_*p*_ *Z*] and 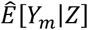 to *E*[*Y*_*m*_|*Z*].

Technically, 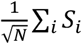 corresponds to a one-step corrected estimator of the expected conditional covariance functional, utilizing the theory of efficient influence functions^5^. Since the power of the test depends on the unknown underlying relationship between *Y*_*m*_ and *X*_*p*_, considering suitable transformations of *X*_*p*_, which are common for omics data and here denoted by *b*_1_(*X*_*p*_), *b*_2_(*X*_*p*_), *…* can improve the performance^6,8^. For a finite set of *J* pre-specified data transformations, indexed by *j*, the individual functionals *E*[*Cov*(*b*_*j*_(*X*_*p*_), *Y*_*m*_ *Z*)] are tested separately, as well as jointly, assuming the number of transformation functions is fixed and small compared to the sample size. Finally, all resulting p-values are also aggregated using the Aggregated Cauchy Association Test (ACAT)^9^.

In general, double machine learning and one-step corrected estimator approaches involve multiple estimation or prediction tasks, followed by plugging the resulting models into test statistics. This is often combined with K-fold cross-fitting^10,11^. Here, the data is split into *K* folds, and the models are trained and plugged into the test statistics in non-overlapping parts, iterating through the folds. Cross-fitting aims to reduce overfitting, can improve efficiency, and allows for establishing asymptotic normality using less restrictive assumptions about the model estimators^3,10^. Because the roles of the data parts are alternated, the full sample size is utilized, but the computational burden increases. For ROMY-CIT, as discussed by Shah and Peters^6^ as well as Niu et al.^12^, cross-fitting is not strictly necessary to ensure valid inference under the null hypothesis. We implemented versions of ROMY-CIT with and without cross-fitting (see Appendix B in the Supplementary Materials).

### ROMY-COV: (Co)variance effects

The second analysis approach considers two different outcomes 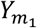 and 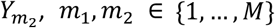, *m*_1_, *m*_2_ ∈ {1, *…, M*} and aims to test the effect of *X*_*p*_ on the covariance between 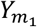 and 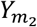. The theoretical analysis of methods for covariance effects attracted increased attention recently, with applications to metabolomics and transcriptomics^13,14^. We define 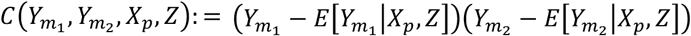 and consider the following functional

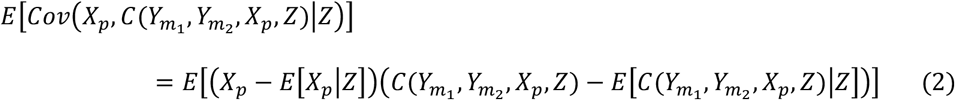

to test the null hypothesis that the covariance between 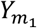 and 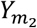 does not depend on *X*_*p*_.

The test statistic is computed by 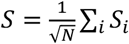 with

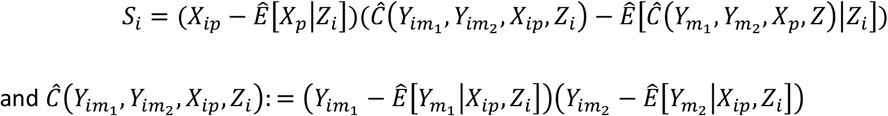

The difference to the mechanics and implementation of ROMY-CIT above is that estimating/training the model 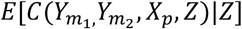 requires plugging in an estimate of 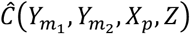 and estimate 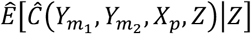 in a second step. In Appendix A of the Supplementary Materials, we derive the efficient influence function for (2) and demonstrate that the one-step corrected estimator corresponds to the proposed test statistic. The algorithm for implementing ROMY-COV is described in Appendix B of the Supplementary Materials. If we set *m*_1_ = *m*_2_, ROMY-COV investigates variance effects.

### ROMY-INTER: Interaction testing

The third analysis component performs interaction-effect testing. We consider the outcome *Y*_*m*_ and implement an approach to test for interaction between *X*_*p*_ and *Z*_*j*_, *j* = 1, *…, D*, in their effect on *Y*_*m*_. Our robust approach, motivated by the work of Vansteelandt et al.^15^, is based on

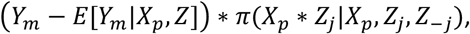

where *π*(*V*|*X*_*p*_, *Z*_*j*_, *Z*_−*j*_) denotes the projection of *V* on functions that have a conditional expectation of 0 given a.) *X*_*p*_ and *Z*_−*j*_ and b.) *Z*_*j*_ and *Z*_−*j*_ (that is, *Z*), where *Z*_−*j*_ denotes the covariates *Z* with *Z*_*j*_ excluded. Based on the derivations of the efficient influence function by Vansteelandt and Dukes^16^, a suitable test statistic is 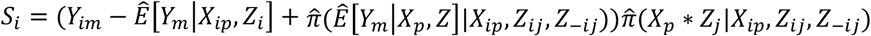. This approach is also related to the approach implemented in the RITSS framework^17^. The intuition is here that *π* performs a projection that provides orthogonality to the main effects of *X*_*p*_ and *Z*_*j*_ and the test statistic is thus less sensitive towards estimation errors of these main effects.

### Statistical learning algorithms

The analysis components require the incorporation of estimators of conditional expectations of the form *E*[*A*|*B*]. These estimators can be realized using different approaches, reflecting assumptions about the complexity of these conditional expectations. We implemented several models, from simple linear regressions to more complex models using generalized additive models (GAMs) and gradient boosting-based approaches such as LightGBM^18^. For the latter two methods, we used the R package *mlr3*^19^. We note that the framework also technically allows for the incorporation of deep learning-based approaches, assuming the sample size provides a reasonable basis for these flexible models.

### Parallel computing

The application of K-fold cross-fitting and flexible statistical learning methods increases the computational burden compared to plain regression analyses. We incorporated the option to utilize parallel computations across all analysis components to ensure the feasibility of large-scale studies. This option uses the machinery of the *BiocParallel* R package, significantly reducing computational time.

### Layer-specific covariates

We implemented the approaches with an option to designate different covariates *Z* to *Y* and *X* for scenarios in which this seems plausible. For example, if domain knowledge can rule out effects on either *Y* or *X*, and the reduced dimensionality might be beneficial for model training/estimation.

## Simulation studies

We conducted numerical experiments to evaluate and illustrate the three analysis components of ROMY. We also included alternative approaches based on standard regressions to highlight the differences and compare the performance.

We implemented a shared simulation setup described in detail in the Supplementary Materials. We considered two different model types for *E*[. |*Z*] and *E*[. |*X, Z*] for the ROMY components. The first is based on a standard linear regression where the conditional mean is a linear combination of the covariates (‘LM’), while the second model approach additionally incorporates the quadratic and product interaction terms for all covariates (‘LMQ’). The corresponding regression-based approach, as an alternative method, is denoted by ‘LR’, and the specification of this regression model depends on the context (see below). The simulations include a scenario in which the covariates *Z* have quadratic effects on *X* and *Y* (‘Q’-scenario), not fully captured by linear effect models. *X* is always a one-dimensional variable and *Z* is two-dimensional in the simulations. All results based on 10,000 replicates, further details are provided in the Supplementary Materials.

### ROMY-CIT

We compared CIT-LM and CIT-LMQ (application of LM and LMQ to *E*[*X*|*Z*] and *E*[*Y*|*Z*] in ROMY-CIT, respectively) with the performance of a linear regression approach based on *E*[*Y*|*X, Z*] = *βX + γ*^*T*^*Z* (robust standard error-based Wald test of *β*, denoted by ‘LR’).

### ROMY-COV

We compared COV-LM and COV-LMQ (application of LM and LMQ to all *E*[· |*Z*] and *E*[· |*X, Z*] in ROMY-COV, respectively) with the performance of a linear regression approach, where we first regressed *Y*_1_ and *Y*_2_ separately on *X* and *Z*. Then we regressed the product of the *Y* residuals on *X* and *Z* and tested the coefficient of *X* (robust standard error-based Wald test, denoted by ‘LR’).

### ROMY-INTER

The potentially interacting variable in all simulations is *Z*_1_. We compared INTER-LM and INTER-LMQ (application of LM and LMQ to *E*[*Y*|*X, Z*] and the ACE algorithm in ROMY-INTER, respectively) with a linear regression approach based on *E*[*Y*|*X, Z*] = *δXZ*_1_ *+ βX + γ*^*T*^*Z* (robust standard error-based Wald test of *δ*, denoted by ‘LR’).

### Results

The simulation studies considered type 1 error rates and power curves. Figures 1-6 visualize the results. Figure 1 shows that LR and CIT-LM have comparable type 1 error rates, which are inflated in scenarios where the confounding effects are not well captured by the conditional expectation models (Q-scenario). CIT-LMQ controls the type 1 error rate in all scenarios due to the model’s ability to account for quadratic effects. Figure 2 shows that the power is similar for all three methods CIT-LMQ, CIT-LM, and LR, showing that the robustness of CIT does not lead to a power loss in this simulation and setup. Figures 3 and 4 show the results of the COV simulations. The results are similar to the CIT observations. The same applies to the INTER results plotted in Figures 5 and 6. However, Figure 6 shows that the linear regression approach LR has slightly higher power, suggesting a trade-off between power and robustness for ROMY-INTER.

**Figure 1.**
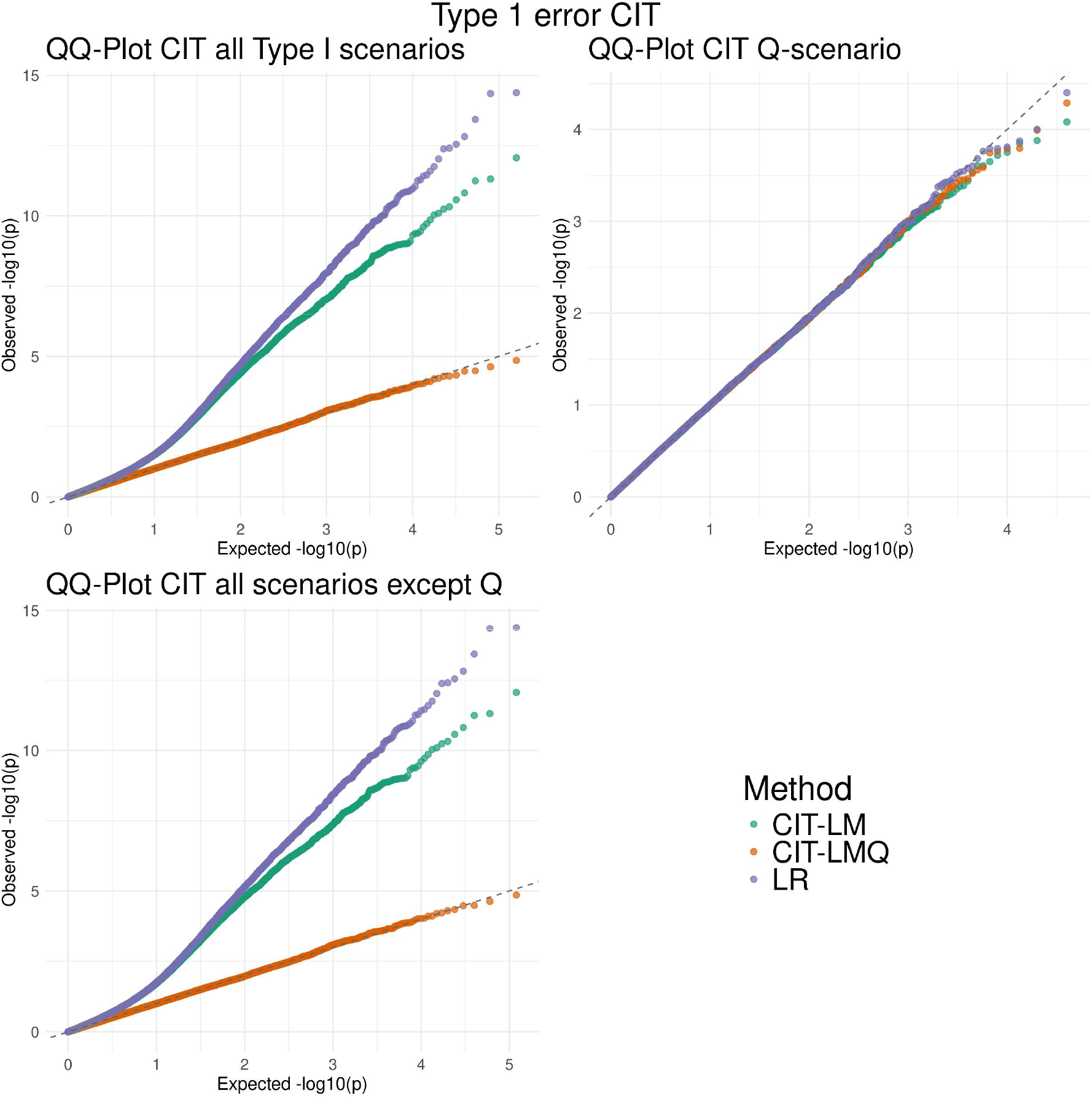
Type 1 error simulations for ROMY-CIT, including a comparison with a standard regression approach ‘LR’.

**Figure 2.**
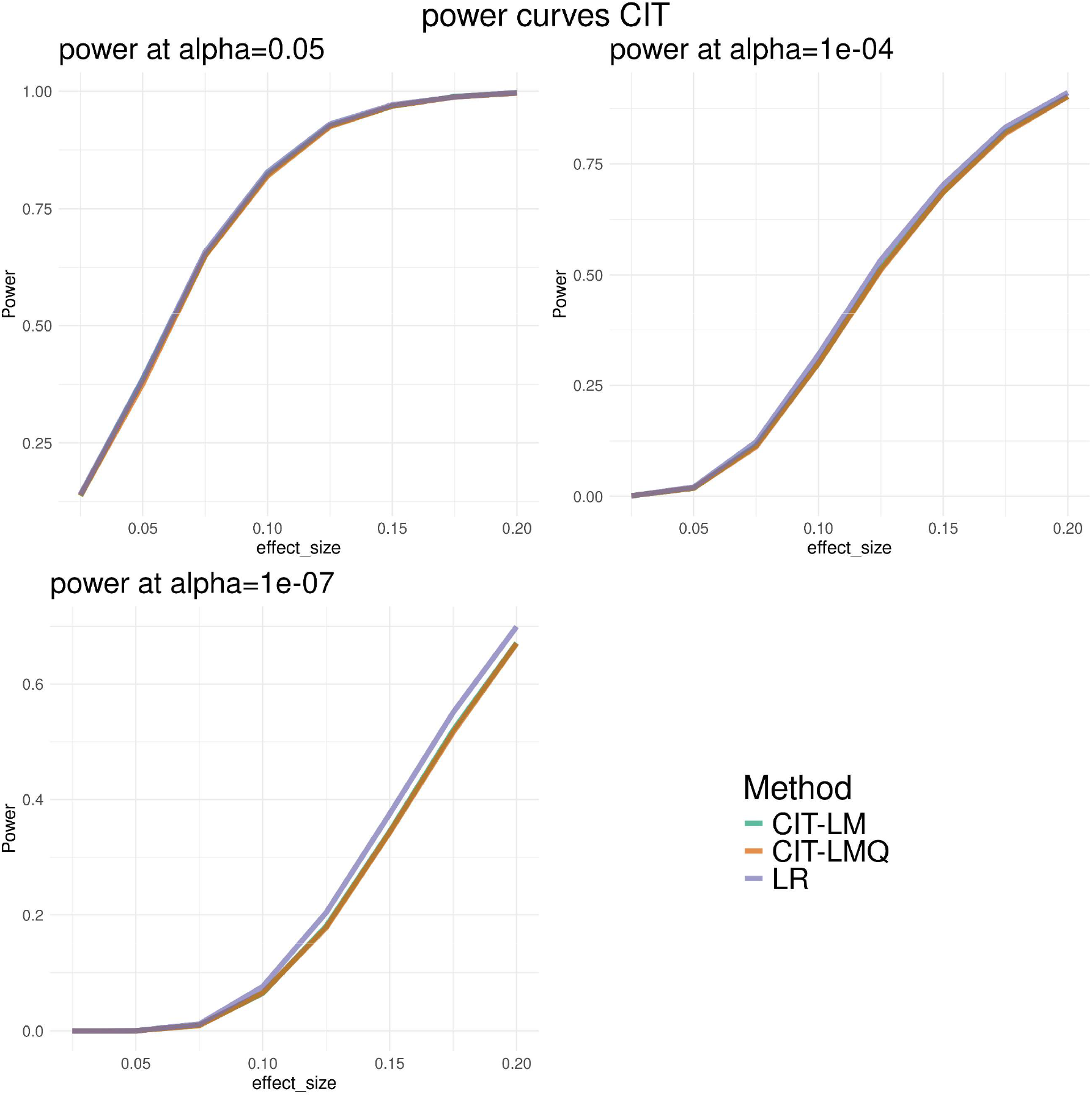
Power simulations for ROMY-CIT, including a comparison with a standard regression approach ‘LR’.

**Figure 3.**
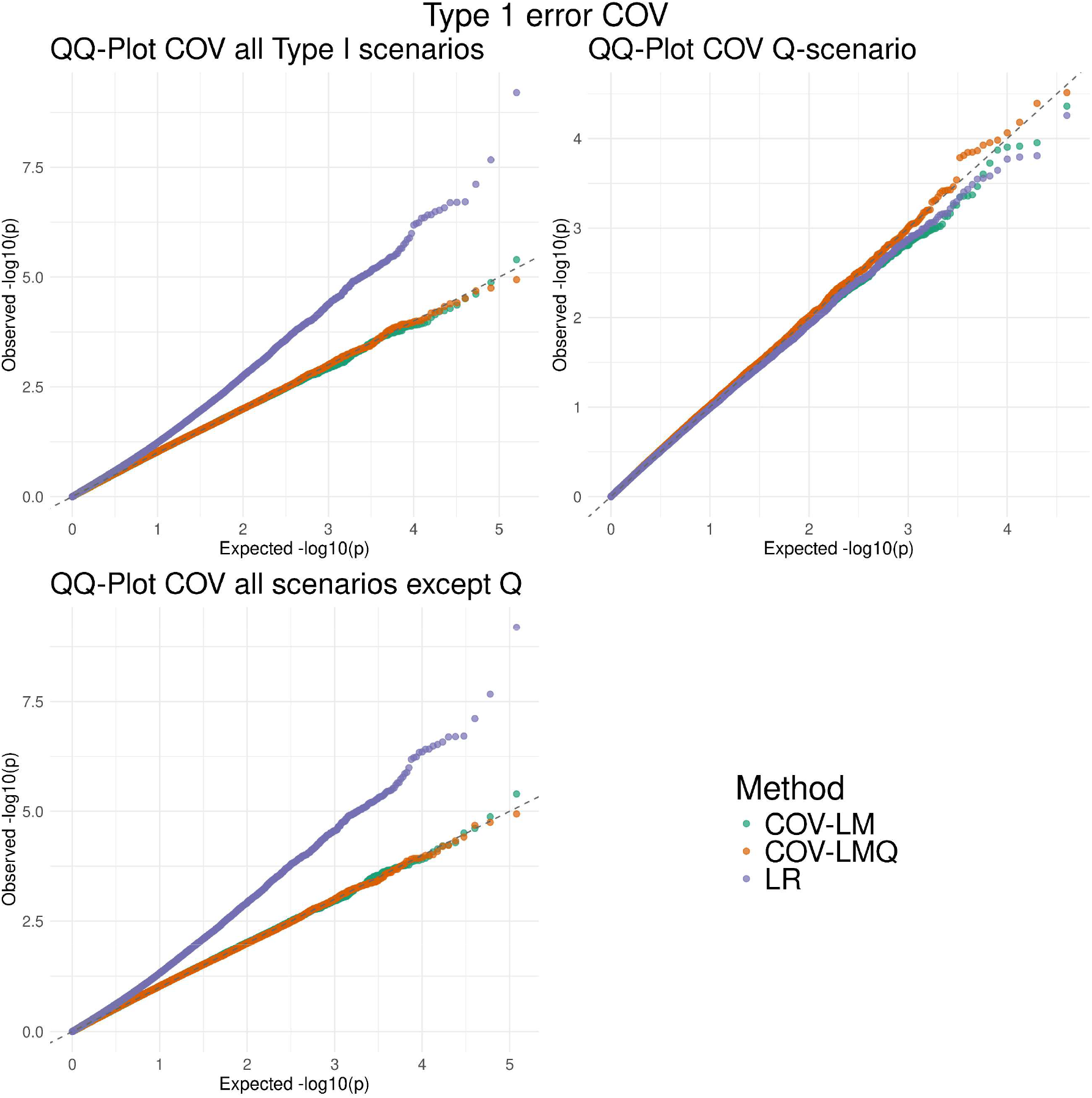
Type 1 error simulations for ROMY-COV, including a comparison with a standard regression approach ‘LR’.

**Figure 4.**
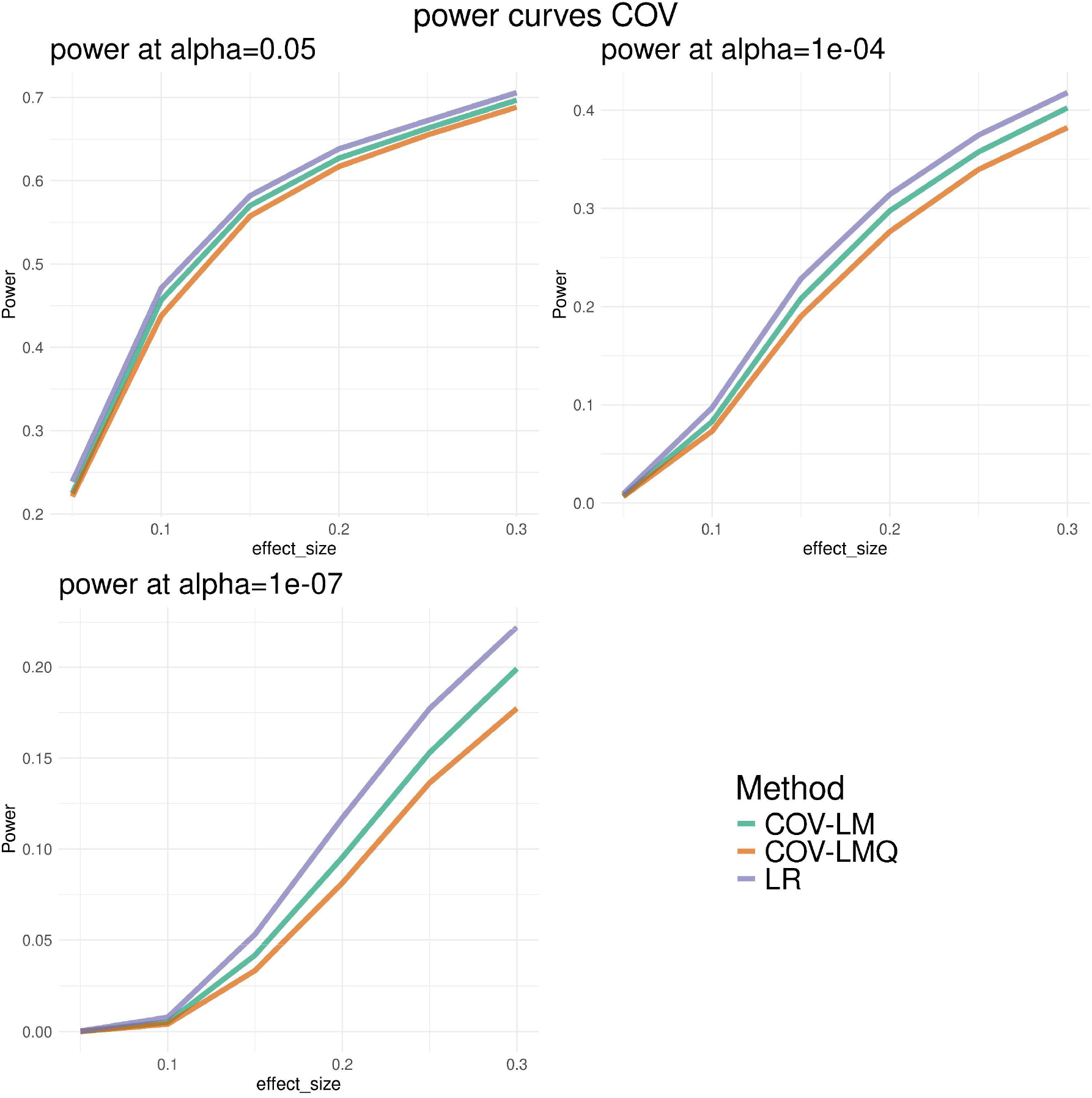
Power simulations for ROMY-COV, including a comparison with a standard regression approach ‘LR’.

**Figure 5.**
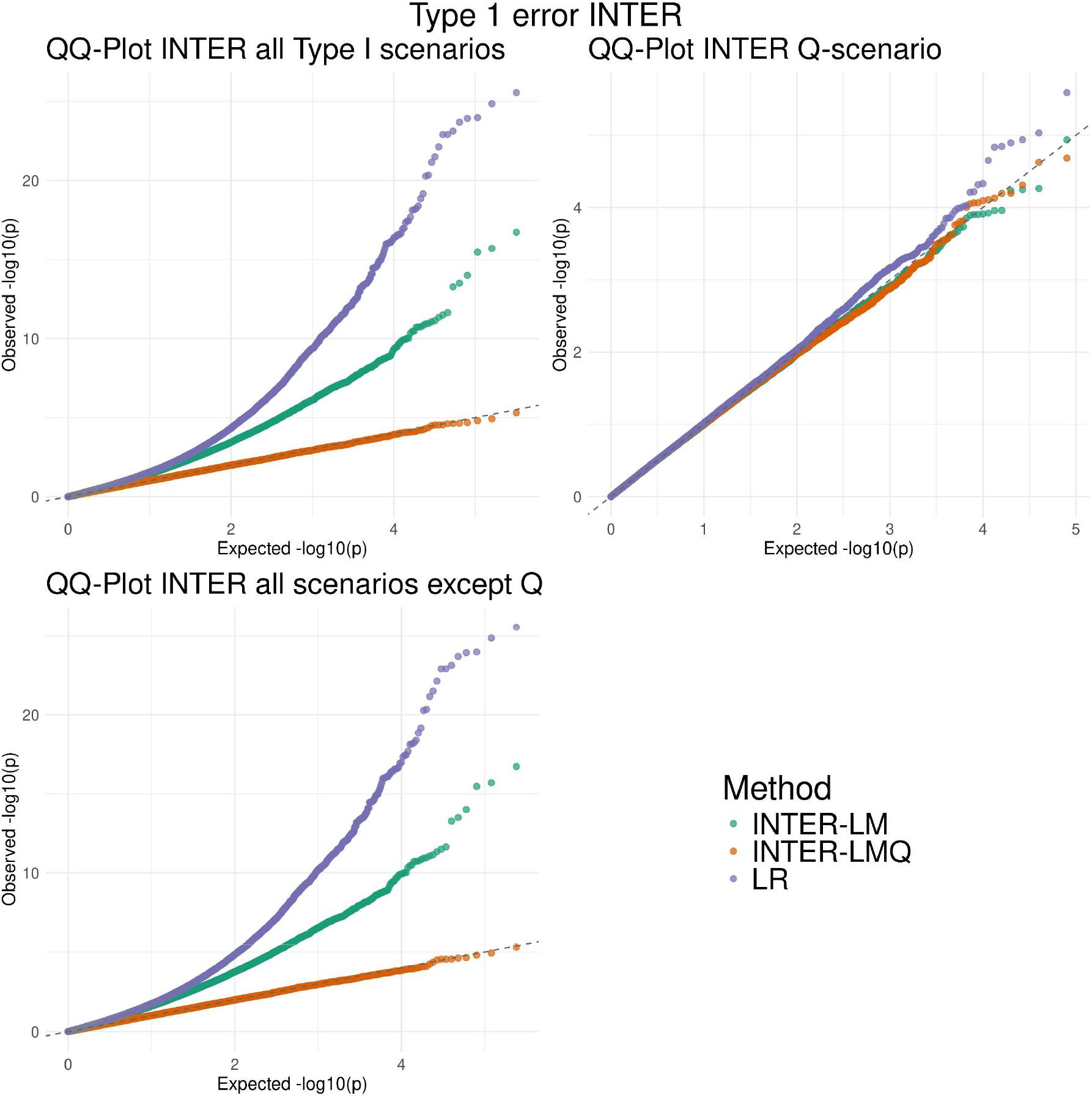
Type 1 error simulations for ROMY-INTER, including a comparison with a standard regression approach ‘LR’.

**Figure 6.**
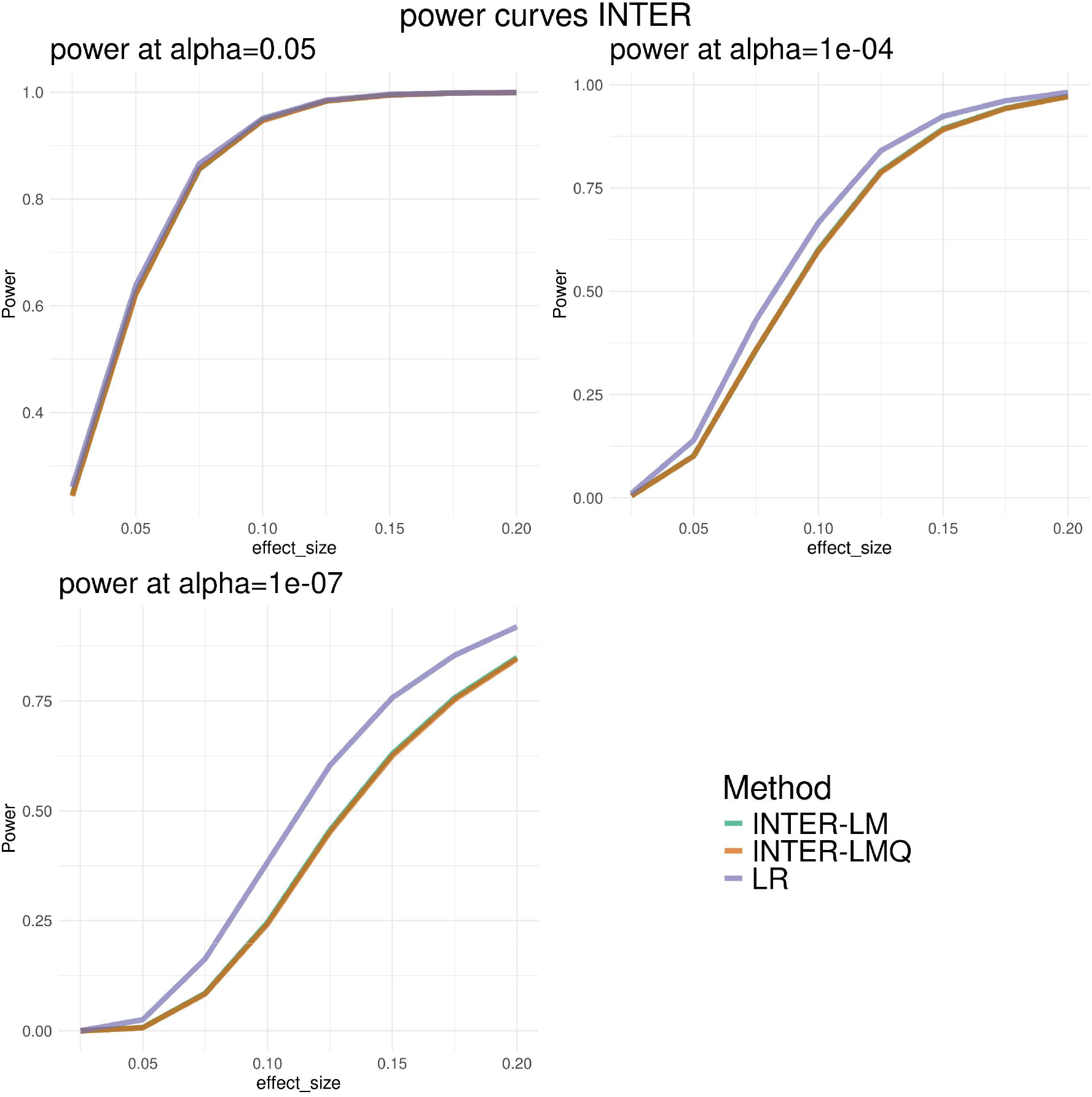
Power simulations for ROMY-INTER, including a comparison with a standard regression approach ‘LR’.

## Discussion

We present our ROMY framework for analyzing multi-omics data. ROMY and the corresponding R package *romy* implement three analysis components that provide flexible, robust methods based on modern advances in double machine learning and theoretical statistics. We illustrated the application of ROMY in a simulation study. The structure of ROMY allows for incorporating future improvements in machine-learning-based prediction models.

