## Supplementary Materials for "Analyzing associations and higher-order effects in multi-omics data with double machine learning"

#### Setting

We denote by $Y = (Y_1, \ldots, Y_M)$ $M$ quantitative outcomes (for example, complex traits or omics measurements). Second, $X = (X_1, \ldots, X_P)$ denotes $P$ factors in a different layer of omics or epidemiological variables. Moreover, $Z$ denotes $D$-dimensional covariates that the analyses aim to adjust for. We assume that this data is collected across a joint set of samples of size $N$, resulting in observed data $Y_i = (Y_{i1}, \ldots, Y_{iM})$, $X_i = (X_{i1}, \ldots, X_{iP})$, and $Z_i = (Z_{i1}, \ldots, Z_{iD})$, $i = 1, \ldots, N$.

#### Appendix A: Derivation of ROMY-COV

In the following, we derive the test statistic to test the effect of $X_p$ on the covariance between $Y_{m_1}$ and $Y_{m_2}$.

##### Test statistic

We will use similar argumentations and notations as Vansteelandt and Dukes (2022). Let $C(Y_1, Y_2, X, Z) := (Y_1 - E[Y_1|X, Z])(Y_2 - E[Y_2|X, Z])$ and define

$$
\begin{aligned}
\Psi &= E[E\left(\{X - E(X|Z)\}\{C(Y_1, Y_2, X, Z) - E[C(Y_1, Y_2, X, Z)|Z]\}|\,Z\right)] \\
&= E\left[\{X - E(X|Z)\}\{C(Y_1, Y_2, X, Z) - E[C(Y_1, Y_2, X, Z)|Z]\}\right] \\
&= \int \{X - E(X|Z)\}\{C(Y_1, Y_2, X, Z) - E[C(Y_1, Y_2, X, Z)|Z]\}\, f(Y_1, Y_2, X, Z) dY_1 dY_2 dX dZ \\
&= \int \{X - E(X|Z)\}\, C(Y_1, Y_2, X, Z) f(Y_1, Y_2, X, Z) dY_1 dY_2 dX dZ \\
&- \int \{X - E(X|Z)\}\, E[C(Y_1, Y_2, X, Z)|Z] f(Y_1, Y_2, X, Z) dY_1 dY_2 dX dZ \\
&= \Psi_1 - \Psi_2
\end{aligned}
$$

We get

$$\begin{aligned}
\frac{\partial \Psi_1}{\partial t}|_{t=0} &= \int \{X - E(X|Z)\} C(Y_1, Y_2, X, Z) S_t(O) f(O) dO \\
&- \int X' S_t(X'|Z) f(X'|Z) C(Y_1, Y_2, X, Z) f(Y_1, Y_2, X, Z) dY_1 dY_2 dX' dX dZ \\
&- \int (X - E(X|Z))(Y_1 - E(Y_1|X, Z)) Y_2' S_t(Y_2'|X, Z) f(Y_2'|X, Z) f(Y_1, Y_2, X, Z) dY_1 dY_2 dY_2' dX dZ \\
&- \int (X - E(X|Z))(Y_2 - E(Y_2|X, Z)) Y_1' S_t(Y_1'|X, Z) f(Y_1'|X, Z) f(Y_1, Y_2, X, Z) dY_1' dY_1 dY_2 dX dZ \\
&= \int \{X - E(X|Z)\} C(Y_1, Y_2, X, Z) S_t(O) f(O) dO \\
&- \int X S_t(X|Z) E[C(Y_1, Y_2, X, Z)|Z] f(X, Z) dX dZ
\end{aligned}$$

and

$$\begin{aligned}
\frac{\partial \Psi_2}{\partial t}|_{t=0} &= \int (X - E(X|Z)) E[C(Y_1, Y_2, X, Z)|Z] S_t(O) f(O) dO \\
&- \int X' S_t(X'|Z) f(X'|Z) E[C(Y_1, Y_2, X, Z)|Z] f(Y_1, Y_2, X, Z) dY_1 dY_2 dX dX' dZ \\
&+ \int (X - E(X|Z)) C(Y_1, Y_2, X', Z) S_t(Y_1, Y_2, X'|Z) f(Y_1, Y_2, X'|Z) f(X, Z) dY_1 dY_2 dX dX' dZ \\
&- \int (X - E(X|Z))(Y_1 - E(Y_1|X', Z)) Y_2' S_t(Y_2'|X', Z) f(Y_2'|X', Z) f(Y_1, Y_2, X'|Z) f(X, Z) dY_1 dY_2 dY_2' dX' dX dZ \\
&- \int (X - E(X|Z))(Y_2 - E(Y_2|X', Z)) Y_1' S_t(Y_1'|X', Z) f(Y_1'|X', Z) f(Y_1, Y_2, X'|Z) f(X, Z) dY_1 dY_1' dY_2 dX' dX dZ \\
&= \int (X - E(X|Z)) E[C(Y_1, Y_2, X, Z)|Z] S_t(O) f(O) dO \\
&- \int X S_t(X|Z) E[C(Y_1, Y_2, X, Z)|Z] f(X, Z) dX dZ.
\end{aligned}$$

Here, $S_x$ and $f$ denote the respective score and density.
Therefore, the efficient influence function is given by (Newey, 1990)

$$(X - E[X|Z])(C(Y_1, Y_2, X, Z) - E[C(Y_1, Y_2, X, Z)|Z]) - \Psi$$

and thus, overall, the one-step corrected estimator of $\Psi$ is

$$(X - E[X|Z])(C(Y_1, Y_2, X, Z) - E[C(Y_1, Y_2, X, Z)|Z])$$

We apply this to $Y_1 = Y_{m_1}$, $Y_2 = Y_{m_2}$, $X = X_p$, and $Z = Z$, and thus consider

$$\frac{1}{\sqrt{N}} S_i = \frac{1}{\sqrt{N}} \sum_{i=1}^{N} (X_{ip} - \hat{E}[X_p|Z_i])(\hat{C}(Y_{im_1}, Y_{im_2}, X_{ip}, Z_i) - \hat{E}[\hat{C}(Y_{m_1}, Y_{m_2}, X_p, Z)|Z_i]),$$

where

$$\hat{C}(Y_{im_1}, Y_{im_2}, X_i, Z_i) = (Y_{im_1} - \hat{E}[Y_{m_1}|X_{ip}, Z_i])(Y_{im_2} - \hat{E}[Y_{m_2}|X_{ip}, Z_i]).$$

#### Asymptotic validity

The asymptotic validity of ROMY-COV can be established with similar arguments as described by Shah and Peters (2020) for the generalized covariance measure (Theorem 6, Shah and Peters (2020)), which corresponds to ROMY-CIT. The difference lies in the fact that ROMY-COV involves the estimator

$$\hat{E}\left[(Y_1 - \hat{E}[Y_1|X, Z])(Y_2 - \hat{E}[Y_2|X, Z])\middle| Z\right]$$

which requires estimating the conditional expectation of an estimated quantity. A discussion of such a scenario can be found in Kennedy (2023). The theoretical conditions to establish asymptotic normality involve standard regularity assumptions, suitable fast product rates of individual convergence rates, and the application of a linear smoother (Kennedy, 2023) model for the second step conditional expectation estimation in

$$\hat{E}\left[(Y_1 - \hat{E}[Y_1|X,Z])(Y_2 - \hat{E}[Y_2|X,Z])\middle|Z\right].$$

Under these assumptions, one can establish asymptotic normality and the consistency of the empirical variance estimator.

### Appendix B: Algorithms

In the following, we describe the algorithms for the three ROMY analysis components.
Each algorithm performs several estimation and plug-in tasks. This can be realized using sample splitting and cross-fitting, or without sample splitting. The sample sets for the estimation and plug-in tasks are denoted by $I_{kj} \subseteq \{1,\ldots,N\}$ and $I_k^c \subseteq \{1,\ldots,N\}$, for $k = 1,\ldots,K$ and $j = 1,\ldots,T$, where $T$ is the number of estimation tasks for the specific algorithm. These sets are generated randomly. If no sample splitting is used, $K = 1$ and $I_1 = I_{1j}^c = \{1,\ldots,N\}$. Otherwise, if $K > 1$, the sample sets of the plug-in and estimation tasks are constructed such that the plug-in set and the combined set of estimation samples do not overlap for fixed $k$.

#### ROMY-CIT

##### Algorithm

The following algorithm describes the association testing step for the variables $Y_m$ and $X_p$, while adjusting for $Z$.

- For each $k$ in $\{1,\ldots,K\}$:
  - Estimate $\hat{E}[b_j(X_p)|Z]$ in $I_{k1}^c$ for all functions $b_j(.), j = 1,\ldots,J$.
  - Estimate $\hat{E}[Y_m|Z]$ in $I_{k2}^c$.
  - Compute the plug-in estimator $S_{ij} = [b_j(X_{ip}) - \hat{E}[b_j(X_p)|Z_i]][Y_{im} - \hat{E}[Y_m|Z_i]]$ in $I_k$.
- Compute combined statistics $S_j = \frac{1}{\sqrt{N}}\sum_{i=1}^N S_{ij}$ and the corresponding empirical covariance matrix.
- Test $S_j$ using standard asymptotic normality theory, separately and jointly (using estimated covariance matrix). Combine the different tests using the ACAT (Liu et al., 2019).

##### Implementation

As discussed by Shah and Peters (2020) as well as Niu et al. (2024), sample-splitting is not needed to ensure asymptotic validity under the null hypothesis, given reasonable regularity conditions. In this case, $K = 1$ and $I_1 = I_{11}^c = I_{12}^c$ ('no-split'). We also implemented sample-splitting options where a.) $K > 1$ and $I_{k1}^c$ and $I_{1k2}^c$ are equal splits of $I_k^c$ ('double-split'), and 2.) $K > 1$ and $I_{k1}^c = I_{k2}^c$ ('single-split').

#### ROMY-COV

##### Algorithm

The following algorithm describes the (co)variance testing step for the variables $Y_{m_1}, Y_{m_2}$, and $X_p$, while adjusting for $Z$. By setting $m_1 = m_2$, this approach can be used to test the effect of $X_p$ on the variance of $Y_{m_1}$, while adjusting for $Z$.

- For each $k \in \{1, \ldots, K\}$:
  - Estimate $\hat{E}[X_p|Z]$ in $I_{k1}^c$
  - Estimate $\hat{E}[Y_{m_1}|X_p, Z]$ in $I_{k2}^c$.
  - Estimate $\hat{E}[Y_{m_2}|X_p, Z]$ in $I_{k3}^c$.
  - Estimate $\hat{E}[\hat{C}(Y_{m_1}, Y_{m_2}, X_p, Z)|Z]$ in $I_{k4}^c$, based on the results from the previous steps that allow to estimate $\hat{C}(Y_{m_1}, Y_{m_2}, X_p, Z)$ by plugging in the estimators.
  - Compute plug-in statistic
    $S_i = [X_{ip} - \hat{E}[X_p|Z_i]][\hat{C}(Y_{im_1}, Y_{im_2}, X_{ip}, Z_i) - \hat{E}[\hat{C}(Y_{m_1}, Y_{m_2}, X_p, Z)|Z_i]]$ in $I_k$.
- Compute combined statistic $S = \frac{1}{\sqrt{N}} \sum_{i=1}^{N} S_i$ and the corresponding empirical variance estimator.
- Test $S$ using standard asymptotic normality theory.

##### Implementation

We implemented the ROMY-COV approach with $K > 1$ and $I_{k1}^c$, $I_{1k2}^c$, $I_{1k3}^c$, and $I_{1k4}^c$ are equal splits of $I_k^c$. We are using a linear smoother for the second stage prediction task (estimating $\hat{E}[\hat{C}(Y_{m_1}, Y_{m_2}, X_p, Z)|Z]$).

#### ROMY-INTER

The following algorithm describes the interaction testing step for the variables $Y_m$ and the interaction between $X_p$ and $Z_j$, while adjusting for main effects of $X_p$ and $Z$.

- For each $k \in \{1, \ldots, K\}$:
  - Estimate $\hat{E}[Y_m|X_p, Z]$ in $I_{k1}^c$.
  - Obtain $\hat{\pi}(X_p * Z_j|X_p, Z)$ objects in $I_{k2}^c$.
  - Obtain $\hat{\pi}(\hat{E}[Y_m|X_p, Z]|X_p, Z)$ objects in $I_{k3}^c$.
  - Compute plug-in estimator $S_i = (Y_{im} - \hat{E}[Y_m|X_{ip}, Z_i] + \hat{\pi}(\hat{E}[Y_m|X_p, Z]|X_{ip}, Z_{ij}, Z_{-ij}))\hat{\pi}(X_p * Z_j|X_{ip}, Z_{ij}, Z_{-ij})$ in $I_k$.
- Compute combined statistic $S = \frac{1}{\sqrt{N}} \sum_{i=1}^{N} S_i$ and the corresponding empirical variance estimator.
- Test $S$ using standard asymptotic normality theory.

##### Implementation

We implemented the ROMY-INTER approach with $K > 1$ and $I_{k1}^c$, $I_{1k2}^c$, and $I_{1k3}^c$ are equal splits of $I_k^c$.

#### Appendix C: Simulation studies

In this section, we describe the simulation studies.

##### ROMY-CIT

###### Setting

- We set $D = 2$ and draw $\mu_Z \sim N(0, I_2)$, where $I_2$ is the 2-dimensional identity matrix, as well as $\sigma_{Z_d} \sim \text{Unif}(1, 2)$, $d = 1, 2$, independently. The covariates $Z_{id}$ are then drawn according to $Z_{id} \sim N(\mu_{Z_d}, \sigma^2_{Z_d})$, for $d = 1, 2$.
- We set $P = 1$ and draw $\mu_X \sim N(0, 1)$, $\sigma_X \sim \text{Unif}(1, 2)$, and $X_i \sim N(\mu_X + f(Z_i), \sigma^2_X)$ for different choices of $f(Z)$.
- Finally, setting $M = 1$, we draw $\mu_Y \sim N(0, 1)$, $\sigma_Y \sim \text{Unif}(1, 2)$, and $Y_i = \mu_Y + g(Z_i) + \beta_X X_i + \delta_{XZ} X_i Z_{i1} + \epsilon_i(X_i)$ for different choices of $g(Z)$, where $\epsilon_i(X_i)$ is a specified distribution with mean 0 and standard deviation $\sigma_Y(1 + \gamma_X|X_i|)$.

In type 1 error and power simulations, all results are based on 10,000 replicates.

###### Type 1 error

We considered the following 16 combinations of simulation parameter specifications for the type 1 error simulations, with $\beta_X = 0$ and $\delta_{XZ} = 0$ fixed:

- $\gamma_X \in \{0.0, 0.1\}$
- $f(Z) \in \{0.1 * Z_1, 0.1 * Z_1^2\}$
- $g(Z) \in \{0.1 * Z_1, 0.1 * Z_1^2\}$
- $\{\epsilon_i \sim N, \epsilon_i \sim \chi^2(1) - 1\}$, where $N$ denotes a normal distribution and $\chi^2(1)$ is a chi-squared distribution with one degree of freedom.

###### Power

We fixed $\delta_{XZ} = 0$, $\gamma_X = 0$, $f(Z) = 0.0$, $g(Z) = 0.0$, and $\epsilon_i \sim N(0, 1)$ and considered

- $\beta_X \in \{0.025, 0.05, 0.075, 0.1, 0.125, 0.15, 0.175, 0.2\}$

###### Applied methods

We applied the ROMY-CIT association testing approach with $K = 5$, $b_1(x) = x$, and two different types of adjustment models to estimate $E[b_1(X)|Z]$ and $E[Y|Z]$. The first model implemented a simple linear regression, the second incorporated higher-order and interaction terms. We denote these two ROMY-CIT implementations as *CIT-LM* and *CIT-LMQ*. We compared these two approaches with a linear regression based on $E[Y|X, Z] = \beta X + \gamma^T Z$ with robust standard error estimation, denoted by *LR*.

#### ROMY-COV

##### Setting

- We set $D = 2$ and draw $\mu_Z \sim N(0, I_2)$, where $I_2$ is the 2-dimensional identity matrix, as well as $\sigma_{Z_d} \sim$ Unif$(1, 2)$, $d = 1, 2$, independently. The covariates $Z_{id}$ are then drawn according to $Z_{id} \sim N(\mu_{Z_d}, \sigma^2_{Z_d})$, for $d = 1, 2$.

- We set $P = 1$ and draw $\mu_X \sim N(0, 1)$, $\sigma_X \sim$ Unif$(1, 2)$, and $X_i \sim N(\mu_X + f(Z_i), \sigma^2_X)$ for different choices of $f(Z)$.

- Finally, setting $M = 2$, we draw $\mu_{Y_m} \sim N(0, 2)$, $\sigma_{Y_m} \sim$ Unif$(1, 2)$, and $Y_{im} = \mu_{Y_m} + g(Z_i) + \beta_X X_i + \delta_{XZ} X_i Z_{i1} + \epsilon_{im}(X_i)$, $m = 1, 2$, for different choices of $g(Z)$, where $\epsilon_i(X_i)$ is a specified distribution with mean 0 and standard deviation $\sigma_{Y_m}(1 + \gamma_X |X_i|)$.

In type 1 error and power simulations, all results are based on 10,000 replicates.

##### Type 1 error

We considered the following 16 combinations of simulation parameter specifications for the type 1 error simulations, with $\delta_{XZ} = 0$ and $\gamma_X = 0$ fixed:

- $\beta_X \in \{0.0, 0.1\}$
- $f(Z) \in \{0.1 * Z_1, 0.1 * Z_1^2\}$
- $g(Z) \in \{0.1 * Z_1, 0.1 * Z_1^2\}$
- $\{\epsilon_i \sim N, \epsilon_i \sim \chi^2(1) - 1\}$, where $N$ denotes a normal distribution and $\chi^2(1)$ is a chi-squared distribution with one degree of freedom.

The type 1 error results focus on the testing of the conditional covariance between $Y_1$ and $Y_2$.

##### Power

We fixed $\delta_{XZ} = 0$, $\beta_X = 0.1$, $f(Z) = 0.0$, $g(Z) = 0.0$, and $\epsilon_i \sim N(0, 1)$ and considered

- $\gamma_X \in \{0.05, 0.1, 0.15, 0.2, 0.25, 0.3\}$

The power results focus on testing the effect on the variance of $Y_1$.

##### Applied methods

We applied the ROMY-COV approach with $K = 5$ and two different types of adjustment models to estimate $E[X|Z]$ and $E[Y_m|X, Z]$, $m = 1, 2$. The first model implemented a simple linear regression, the second incorporated higher-order and interaction terms. For both approaches, we applied simple linear regression to perform the estimation of $E[C(Y_1, Y_2|X, Z)|Z]$. We denote the two ROMY-COV implementations as *COV-LM* and *COV-LMQ*. We compared these two approaches with a linear regression based on $E[Y_m|X, Z] = \beta_m X + \gamma_m^T Z$ to obtain the product of these two residuals and perform a second step regression of this product on $X$ and $Z$, with robust standard error estimation, denoted by *LR*.

##### ROMY-INTER

###### Setting

- We set $D = 2$ and draw $\mu_Z \sim N(0, I_2)$, where $I_2$ is the 2-dimensional identity matrix, as well as $\sigma_{Z_d} \sim \text{Unif}(1, 2)$, independently. The covariates $Z_{id}$ are then drawn according to $Z_{id} \sim N(\mu_{Z_d}, \sigma^2_{Z_d})$, for $d = 1, 2$.

- We set $P = 1$ and draw $\mu_X \sim N(0, 1)$, $\sigma_X \sim \text{Unif}(1, 2)$, and $X_i \sim N(\mu_X + f(Z_i), \sigma^2_X)$ for different choices of $f(Z)$.

- Finally, setting $M = 1$, we draw $\mu_Y \sim N(0, 1)$, $\sigma_Y \sim \text{Unif}(1, 2)$, and $Y_i = \mu_Y + g(Z_i) + \beta_X X_i + \delta_{XZ} X_i Z_{i1} + \epsilon_i(X_i)$ for different choices of $g(Z)$, where $\epsilon_i(X_i)$ is a specified distribution with mean 0 and standard deviation $\sigma_Y(1 + \gamma_X|X_i|)$.

In type 1 error and power simulations, all results are based on 10,000 replicates.

###### Type 1 error

We considered the following 32 combinations of simulation parameter specifications for the type 1 error simulations, with $\delta_{XZ} = 0$ fixed:

- $\beta_X \in \{0.0, 0.1\}$
- $\gamma_X \in \{0.0, 0.1\}$
- $f(Z) \in \{0.1 * Z_1, 0.1 * Z_1^2\}$
- $g(Z) \in \{0.1 * Z_1, 0.1 * Z_1^2\}$
- $\{\epsilon_i \sim N, \epsilon_i \sim \chi^2(1) - 1\}$, where $N$ denotes a normal distribution and $\chi^2(1)$ is a chi-squared distribution with one degree of freedom.

###### Power

We fixed $\beta_X = 0.1$, $\gamma_X = 0$, $f(Z) = 0.0$, $g(Z) = 0.0$, and $\epsilon_i \sim N(0, 1)$ and considered

- $\delta_{XZ} \in \{0.025, 0.05, 0.075, 0.1, 0.125, 0.15, 0.175, 0.2\}$

###### Applied methods

We applied ROMY-INTER with $K = 5$ and two different types of adjustment models to estimate $E[X|Z]$, $E[Y|X, Z]$ and the ACE algorithm adjustments. The first model implemented a simple linear regression, the second incorporated higher-order and interaction terms. We denote the two ROMY-INTER implementations as *INTER-LM* and *INTER-LMQ*. We compared these two approaches with a linear regression based on $E[Y|X, Z] = \delta X Z_1 + \beta X + \gamma^T Z$ with robust standard error estimation, denoted by *LR*.
